# Captamer: A Novel Quantitative Protein Detecting Method Depending on Aptamer-activated Molecular Switches and RPA Signal Amplification

**DOI:** 10.1101/2025.11.02.686180

**Authors:** Yaozhong Cao, Mai Li, Guangtao Xu, Shuyan Xia, Xiaoyu Wu, Keyue Shi, Ruihao Xue, Han Wang, Rundong Ye, Zijun Han, Jinqiang Xu, Qian Zhang, Jiong Hong

**Affiliations:** National Training Center for Laboratory Techniques of Life Science, Department of Life Sciences and Medicine, University of Science and Technology of China. No. 96 Jinzhai Road, Shushan District, Hefei City, Anhui Province 230026, China

**Author notes:** Corresponding Author Jiong Hong- Department of Life Sciences and Medicine, University of Science and Technology of China., Qian Zhang- Department of Life Sciences and Medicine, University of Science and Technology of China. These authors contributed equally to this work. Yaozhong Cao, Mai Li, and Guangtao Xu prepared the figures and wrote the manuscript. Yaozhong Cao, Guangtao Xu, and Rundong Ye designed the experiment and analyzed the data. Mai Li, Shuyan Xia, and Xiaoyu Wu completed the main experimental work. Qian Zhang and Ruihao Xue conducted fund management. Keyue Shi provides ideas for scheme mapping. Keyue Shi, Jinqiang Xu, han Wang, and Zijun Han contributed to the research and pre-experiments. Jiong Hong and Qian Zhang supervised the project. All authors have given approval to the final version of the manuscript.

**Keywords:** Protein assay, Aptasensor, Recombinase polymerase amplification, SARS-CoV-2 nucleocapsid protein, Tau protein, Thrombin

## Abstract

There are various protein assays for specific and quantitative detection and widely used for laboratory and clinic purposes, but current methods still have limitations. Immunoassays based on antibodies, like ELISA, suffer from slow response and a long antibody-screening period, while physical or electrochemical methods are generally restricted by high cost or the stringent requirement of equipment or operating skills. In this study, we developed an in vitro sensitive protein quantification method: Captamer. The Captamer system comprises a molecular switch derived from aptamer sequence and an exponential fluorescence signal amplification pathway based on recombinase polymerase amplification (RPA). We demonstrated the Captamer for SARS-CoV-2 nucleocapsid protein detection and obtained results from samples within 30 min, displaying a wide detection window from 0.2 pg/mL to 200 pg/mL with high specificity. Furthermore, we tested the Captamer for Tau441 protein (a potential Alzheimer’s disease biomarker) and thrombin (a classic aptamer-protein interaction model), showing the limit of detection as low as 1 ng/mL and 0.02pg/mL respectively, which suggested the capacity of Captamer to be applied to various aptamer-protein pairs. Compared with the most commonly used and recently reported protein quantification methods, Captamer stands out for its high sensitivity, short response time, low cost, and simplicity, indicating its great potential to be widely used in protein quantification.

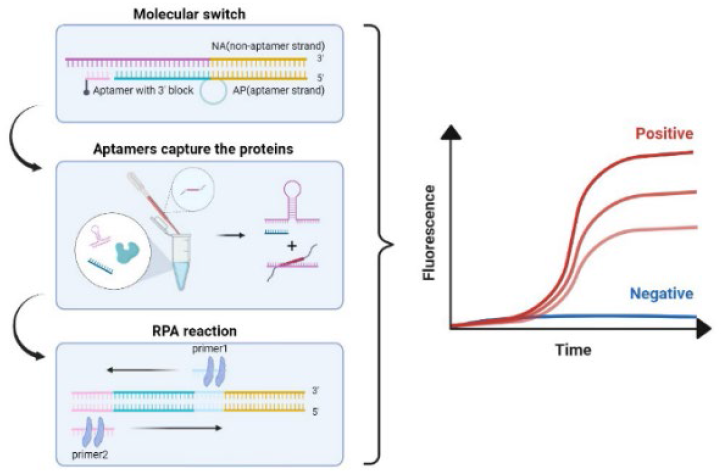

---

*In vitro* specific protein quantification is an essential biochemical technology widely used in biological, medical, environmental, and industrial analyses. The general advantages and limitations of several most commonly used methods were briefly summarized in Table 1. Immunoassays based on antibodies, like ELISA (enzyme-linked immunosorbent assay), are the most popular among these methods because of their accuracy, robustness, versatility, and low cost, but they still have limitations, such as slow response and a long antibody-screening period^1–5^. With the catcher molecules (antibody or aptamer) immobilized on the surface of chips or electrodes, physical and electrochemical biosensors transform the target protein concentration to the change of liquid-solid surface properties which can be detected easier and faster, so they generally have rapid response and high sensitivity. However, they are prone to high cost or requirement of special devices, which prevent their wide replacement of traditional immunoassays ^6–8^. Another strategy is to transform protein level to DNA level. Proteins lack effective amplification methods, while DNA can be exponentially amplified by PCR. Therefore, PCR-based DNA quantification methods have several orders of magnitudes higher sensitivity and wider detection window than most traditional immunoassays ^9^ . By coupling DNA ligation with the specific binding between the target protein and the catcher molecules, PLA (proximity ligation assay) transforms detecting protein into detecting DNA nucleic acid sequence by real-time PCR. Thus, it performs excellently in specific and quantitative protein detection, manifesting higher sensitivity than traditional immunoassays^10^.

**Table 1.** General advantages and limitations of the most commonly used specific protein assays.

| Methods | Principles | Advantages | Limitations |
| --- | --- | --- | --- |
| ELISA<sup>1</sup> | Antibody | Low-cost | Complex steps; Cross-reactivity |
| WB<sup>2</sup> | Electrophoresis; Antibody | Ultra-specific; Robust; Low-cost | Complex steps |
| LFA<sup>22</sup> | Antibody /aptamer | Portable; Rapid; Low-cost; Simple | Precision; Sensitivity |
| PLA<sup>10</sup> | Antibody /aptamer;<br>Real-time PCR | Sensitive; Versatile | Two steps reaction;<br>Requirement of two epitopes |
| Affinity biosensors<sup>8</sup> | Antibody /aptamer;<br>Physical or electrochemical property sensors | Rapid-response; Sensitive;<br>High-throughput;<br>Readily-miniaturized | Cost;<br>Stability;<br>Device;<br>Resistance to noise |
| MS<sup>23</sup> | Mass-to-charge ratio | Ultra-specific; Multiplexing;<br>Free of antibody/aptamer | Sensitivity; Device;<br>Sample preparation |

Aptamers are single-stranded oligonucleotides that can specifically bind to target molecules, particularly proteins^11^. As alternative catcher molecules to antibodies, aptamers exhibit advantages in terms of storage stability, synthesis cost, and screening period^12,13^. Most importantly, they are much easier to be linked with DNA amplification. Based on the operating principle and downstream signal output, aptamer sensors can be broadly classified into conventional types that rely on DNA output signals ^14–16^, antibody-dependent types that couple with antibodies^17,18^, and electrochemical sensors that rely on physicochemical signals such as current value^19–21^.

In the conventionals types of aptasensors, the aptamer is generallyused as an initiator/repressor switch, which turns on/off the downstream DNA amplification like PCR via its binding variant with the target protein^14^. However, PCR reactions lead to protein denaturation and require a complicated temperature cycle, so now these aptasensors prefer amplification by means of isothermal amplification such as Loop-Mediated Isothermal Amplification(LAMP) ^24,25^, Catalytic Hairpin Assembly (CHA)^26^ and Entropy-driven Strand Displacement Reaction (ESDR)^27^. RPA is another promising means of isothermal amplification, requiring only 20-40 minutes at 37°C and supporting the target protein’s correct conformation and the aptamer’s binding affinity ^28^. Due to its fast speed and high sensitivity, RPA is now commonly used to detect various pathogenic DNA/RNA and nucleic acid-based biomarkers^29^. Still, few studies have used RPA to detect proteins^17^.

The coronavirus disease 2019 (COVID-19) pandemic caused by severe acute respiratory coronavirus 2 (SARS-CoV-2) has struck most countries in the world, leading to ∼767 million cases and ∼7 million deaths (World Health Organization, 30 May 2023). The prevalent clinically diagnostic method is PCR-based nucleic acid testing (NAT) ^30^. However, NAT needs complex operations and takes hours for detection, while a previous study demonstrated its diagnostic sensitivity as low as 81% ^31^. In comparison, the nucleocapsid protein (NP) of SARS-CoV-2 is a better biomarker for early diagnosis than its genome nucleic acid ^32,33^. Thus, there is an urgent need for a commercial SARS-CoV-2 NP detection kit with high sensitivity and rapid response for COVID-19 diagnosis.

In this study, we developed an *in vitro* specific and quantitative protein detecting method, termed Captamer, which combines the words “capture” and “aptamer”. Captamer comprises two parts, an aptamer-derived DNA molecular switch and a fluorescence signal amplification pathway based on real-time RPA (recombinase polymerase amplification). Here, we demonstrated the Captamer for NP detection and checked its specificity with another three proteins, Tau441 protein (Tau), thrombin (Tb) and hemoglobin (Hb). Tau441 protein is the longest isoform of human microtubule-associated protein tau. As one representative component of cerebrospinal fluid (CSF) total tau, Tau441 protein is regarded as a CSF and blood biomarker in Alzheimer’s disease^34^. Thrombin is a multifunctional serine protease and a classic aptamer-protein interaction model^35^. Hemoglobin is one of the main components of blood protein. Tau441 and thrombin were also tested by Captamer to illustrate the detectable protein diversity of our system.

## MATERIALS AND METHODS

### Design of Captamer

There are three groups of components in the Captamer system: DNA solutions, RPA reagents, and the protein sample to be measured. The previously synthesized ssDNA components include the NA strand (non-aptamer strand), the AP strand (aptamer strand), the aptamer, primer 1, primer 2, and a fluorescence probe (Fig. 1). The aptamer and the probe have a 3’-end block group (C3-spacer here) to eliminate their priming activity (Fig. 1A). The NA strand, the AP strand, and the aptamer together form a molecular-switch-off DNA template in the absence of the target protein, which lacks the binding sequences (complementary to the corresponding oligonucleotides) of primer 1, primer 2, and the probe, so it cannot be amplified by RPA (Fig. 1A) .

**Fig. 1.**
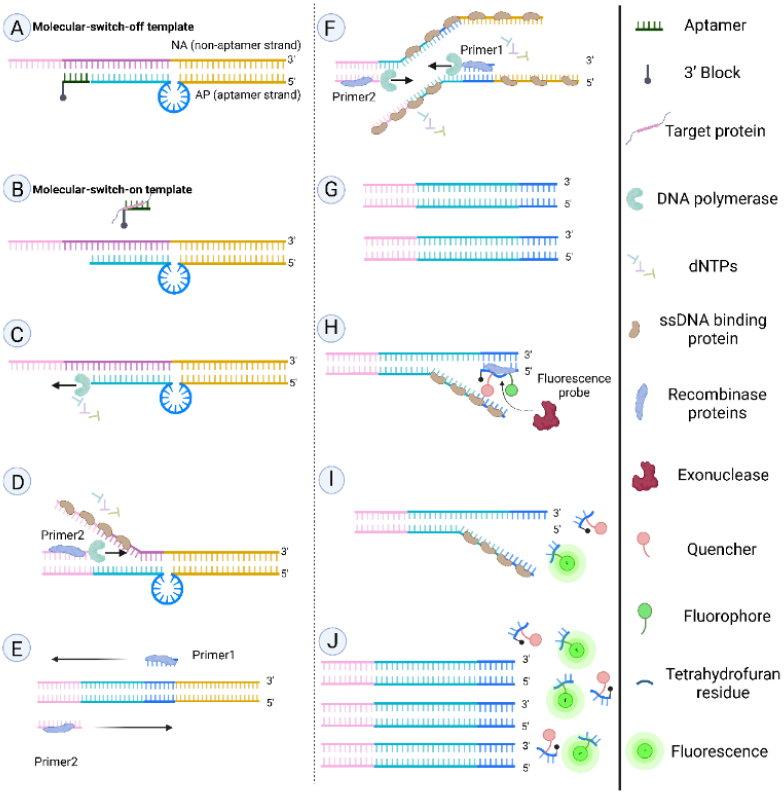
Schematic representation of Captamer. A-B shows the target protein binds to the aptamer to turn on the molecular switch. C-G shows how RPA reaction multiplies molecular-switch-on DNA templates. H-J shows the specific probe cleavage transforms the level of the DNA templates to fluorescence intensity. Created by Biorender.

As shown in Fig. 1, if the target protein exists in the protein sample, it will compete with the NA strand to bind to the aptamer, expose the 3’-end of the AP strand to the DNA polymerase, and initiate the fill-in of 5’-overhang at 3’-terminal of AP strand (Fig. 1.B). Therefore, primer binding sites will appear and start the RPA amplification reaction. (Fig.1.C-G). Then, the probe will recognize the complementary sequence and bind to it, and the nucleic acid exonuclease will cleave the THF site, leading to released fluophores and fluorescent signal (Fig. 1J).

### DNA solution preparation

The full-length 58nt aptamer discovered by Zhang et al. in 2020 ^36^ was called Nor-Apt (normal aptamer) here, which was used for NP detection. The N9 aptamer was generated by truncating the 5’ end of Nor-aptamer by 9 nt. Similarly, the 29 nt core region as the aptamer for Tau was designed through truncating both 5’ 24 nt and 3’ 23 nt flanking regions of the 76 nt full-length aptamer ^34^. The 29 nt aptamer of thrombin^35^ was short enough and thus directly used. Asymmetric PCR was conducted to get ssDNA with an adjusted forward/reverse primer ratio of 20:1. We prepared the storage solution of the DNA molecular switch via DNA hybridization by mixing the three DNA components (NA strand: AP strand was 1:1, and the aptamer concentrations varied at different experiment setups, ranging from 0-3 times the NA concentration) and renaturing it from 95°C to 28°C at the rate of -6 °C/5 min in a thermocycler for 1h. We stored the storage solution at 4°C and diluted it to the target concentration before usage. Detailed protocol of DNA preparation and sequences were concluded in the Supplementary Materials.

### Real-time RPA assay

Real-time RPA assays were conducted with the DNA Thermostat Rapid Fluorescence Kit (Nanjing Warbio Biotechnology Co., Ltd) according to its manufacturer’s manual. The initial DNA components in a real-time RPA system were called one set of DNA for real-time RPA later, including a pair of primers, a probe, and a molecular switch. The mock group referred to the real-time RPA system in which ddH_2_O replaced the molecular switch. The positive control was molecular switch with no aptamer to measure the highest fluorescence intensity in captamer system.

### Optimization of RPA reaction for molecular switch

Two sets of DNA were designed including NA/AP, primer1/2 and probe. Set1 NA/AP sequence was derived from previous work used for NP and thrombin, while Set2 NA/AP was designed in the basis of GFP sequence for Tau. We mainly used Set1 for the subsequent experiments.

Positive control RPA assays (No aptamer) were conducted at 35°C, 37°C, 39°C, and 41°C (Fig. 2A) to explore the best reaction temperature. After the optimal temperature was determined, the RPA assays were conducted with the aptamer at concentration 0x, 0.5x, 1x, 2x, and 3x of NA strand respectively to select the most suitable aptamer concentration for the switch. Here, we introduced a new concept, block efficiency (BE), to quantitatively characterize the signal-to-noise ratio property of the molecular switch.

**Fig. 2.**
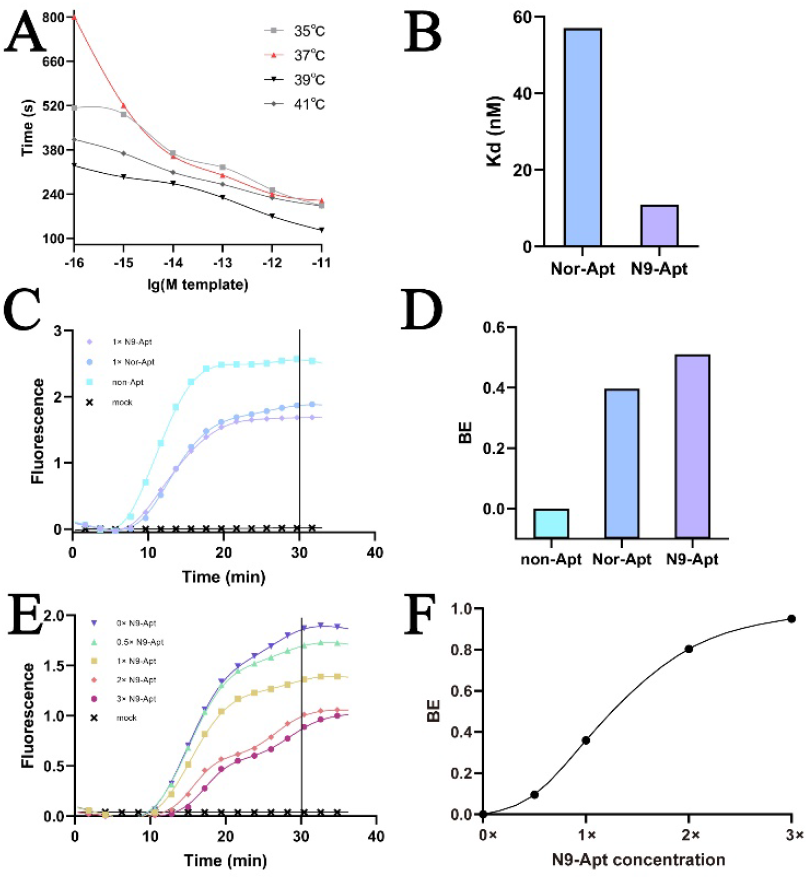
Optimization of the Captamer for NP. **A)** Optimal Temperature Exploration. The time refers the reaction time when the fluorescence intensity values reached 0.2 (data from Fig. S3. A-D). **B)** The dissociation constants between NP and the two aptamers measured by MST. **C)** Real-time RPA assay to compare the molecular switches of N9-Apt and Nor-Apt. **D)** The block efficiency of N9-Apt and Nor-Apt (data from C). **E)** Real-time RPA assay with N9-Apt at different concentrations. **F)** Analysis of BE values of different concentrations of N9-Apt. (data from E).

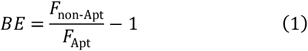

Equation (1) shows the definition of BE of an aptamer at a specific concentration, where *F*_non-Apt_ and *F*_Apt_ refer to the fluorescence intensity value of the group without aptamer (no Apt group) or with the aptamer mentioned above (Apt group) at 30 min in real-time RPA assay. According to this definition, the BE of non-Apt group is zero.

Three batches of real-time RPA assays were conducted to characterize the molecular switch for NP. The first batch compared the BE of 1× Nor-Apt and 1× N9-Apt (“1×” means 1-fold of the molecular switch concentration). The second batch compared the BE of 1× N9-Apt at 37°C and 39°C. The third batch characterized the molecular switch of N9-Apt with various concentrations of N9-Apt (0×, 0.5×, 1×, 2×, and 3×) by comparing their BE.

### Microscale thermophoresis

The affinity of NP to Nor-Apt and N9-Apt was determined by MST (microscale thermophoresis) on Monolith NT.115. NP was labeled with fluorescence using Monolith His-Tag Labeling Kit RED-tris-NTA 2nd Generation (Nano Temper, China). As the dissociation constant (K_d_) between the RED-tris-NTA 2nd Generation dye and NP was stronger than 10 nM, the NP was stained with the dye at twice the concentration of NP. Then, we loaded the Monolith NT.115 Premium Capillary and measured the samples at 40% excitation power and medium MST power. Finally, we obtained the K_d_ with the analysis tool in the Monolith Software.

### Captamer assay

The protocol of the Captamer assay was derived from the real-time RPA assay. Two microliter molecular switch solution (containing 2× aptamer) and 11.5 μL protein sample were mixed at room temperature to prepare 13.5 μL premix as the template solution in a real-time RPA system. The subsequent steps were the same as the real-time RPA assay. Set1 DNA was used for NP and thrombin assays, while Set2 was used for Tau.

### Specificity verification of the Captamer for NP

We conducted another batch of Captamer assay with the NP-Captamer to detect 0.2 ng/mL Tau, 0.1 ng/mL thrombin, and 0.1 ng/mL Hb for specificity verification. The protein sample was replaced with ddH_2_O in the blank group.

We introduced a new value, ΔRelative fluorescence, to integrate and compare the results of Captamer assays from different batches or qPCR machines, which was defined as the result of the protein sample divided by the result of the corresponding blank group minus one. According to this definition, the ΔRelative fluorescence of the mockgroup was zero.

### ELISA assay

We used the ELISA data as a standard to measure the LOD of the captamer assay. The kit manufacturers including 2019-nCoV Nucleocapsid Protein ELISA Kit(Beyotime, China), Human MAPτ(Microtubule Associated Protein Tau/Tau Protein) ELISA Kit(ElabScience, China), human Thrombin ELISA ab270210(Abcam, England).

## RESULTS

In Captamer, the RPA signal amplification pathway transforms the concentration of the molecular-switch-on DNA template into fluorescence intensity. The set of DNA sequences of real-time RPA and the reaction temperature are the two main manually-controlled factors that affect RPA signal amplification, which were explored as follows.

Then, Set1 RPA signal amplification was characterized at 35°C, 37°C, 39°C, and 41°C (Fig. 2. A, Fig. S3. A-D). The curve of the signal amplification at 37°C showed both a steep slope and a wide effective input range from 10^-16^M to 10^-12^M, so 37°C was considered as the best reaction temperature rather than 39°C which is the standard reaction temperature described in the protocol of the real-time RPA kit. Besides, the RPA reaction with lower DNA template concentration was more vulnerable to possible aerosol nucleic acid contamination. Thus, the right margin 10^-12^ M was chosen as the initial DNA molecular switch concentration in Captamer.

According to the design of Captamer, the target protein should compete with the NA strand to bind with the aptamer and activate the molecular switch, so the aptamer prefers a lower affinity to the NA strand and a higher affinity to the target protein for higher sensitivity. As the truncated version of Nor-Apt, N9-Apt is supposed theoretically to bind weaker to the NA strand. We further verified that truncating the 5’ flanking region of Nor-Apt did not reduce its affinity to NP but improved it by more than five times (Fig. 2 B and Fig. S4). Interestingly, the weak binding of the 5’-end is more likely to promote strand displacement than other regions in a DNA oligo^37^. Thus, exposing the core region of Nor-Apt might increase the probability of polymerase-driven strand displacement once the core region bound to NP, which would improve the sensitivity of this molecular switch. Besides, N9-Apt exhibited a slightly higher BE than Nor-Apt (Fig. 2 C, D), which would contribute to lower background noise. We also compared the BE of N9-Apt at 37°C and 39°C (Fig. S3. G), and higher temperature led to lower BE as expected. Therefore, we used the DNA molecular switch of N9-Apt in the Captamer for NP and 37°C is the most suitable for Captamer assay.

Next, we conducted a thorough characterization of the molecular switch derived from N9-Apt and determined the optimal initial aptamer concentration based on its sensitivity and binding efficiency (Fig. 2E, F). As illustrated in Figure. 2 F, a steeper slope indicates a higher sensitivity of the system. Notably, the 2× aptamer concentration exhibited the desired attributes of both a high slope and a high binding efficiency value. Consequently, we established the initial aptamer concentration in the DNA molecular switch as 2×10-^12^ M, representing a two-fold increase compared to the NP concentration.

The Captamer assays were carried out with NP samples at different concentrations (Fig. 3 A). Within 20 min, the fluorescence intensities formed monotonic increasing gradients as the NP concentration increased, so we set 20 min as the endpoint of the Captamer assays and read the fluorescence intensity values at 20 min as the results. All the four NP samples group had results significantly higher than the 0 NP group. They were also significantly different from each other (with fluorescence intensity 1.25, 1.42, 1.53, and 1.69), indicating that the Captamer for NP could detect NP at a concentration as low as 0.2 pg/mL. Then, there was a strong linear correlation with an R-squared of 0.994 between the results and the logarithm of NP concentrations from 0.2 pg/mL to 200 pg/mL (Fig. 3. B). It means that the Captamer for NP had a wide but accurate quantitative detection window.

**Fig. 3.**
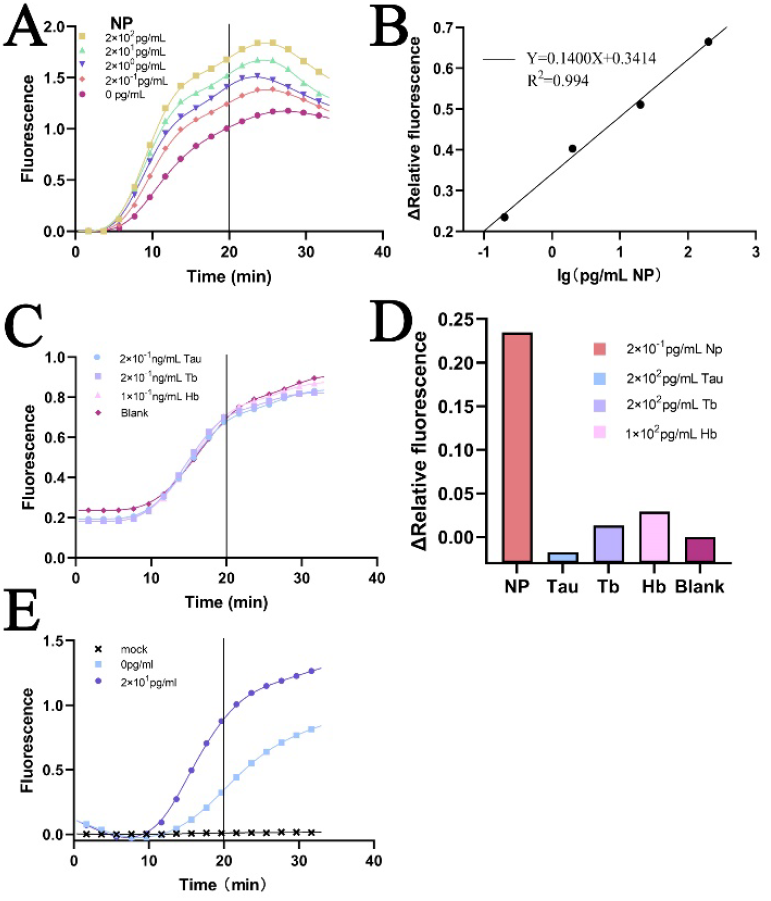
Captamer assays for NP. **A)** Captamer assay for NP. **A)** Linear regression fit between the results of the Captamer assay for NP and the logarithm of the NP concentrations (data from A). **C)** Captamer assay for Tau, Tb, Hb, and ddH_2_O (blank) samples with the molecular switch for NP. **D)** Specificity of the Captamer for NP (data from A and C).

In the actual testing process, it is easier and more convenient to take saliva than blood. Thus, we chose human saliva samples to evaluate the Captamer system’s ability in real biological samples. The results showed that the Captamer system worked well in human saliva. 10 pg/mL of NP had a ΔRelative fluorescence as high as 0.6, which boded well for clinical application (Fig. 3. E-F). To verify the specificity of Captamer, three other proteins were tested with the Captamer for NP, including 0.2 ng/mL Tau, 0.1 ng/mL Tb, and 0.1 ng/mL Hb (Fig. 3C). The real-time RPA curves of these three proteins were almost coincident with the one of the blank groups, indicating the Captamer for NP responded little to Tau, Tb, and Hb. This conclusion was further validated when we calculated the ΔRelative fluorescence of these proteins, which were almost zero, much smaller than 0.23 for NP (Fig. 3D). Even at 500-or 1,000-times lower concentration than the other three proteins, 0.2 pg/mL NP had a significantly higher ΔRelative fluorescence. It suggested the potential application of Captamer to detect biological samples containing complicated components.

To demonstrate the detectable protein diversity of the Captamer system, we replaced the NP aptamer with a new one for Tb detection in water. Just like NP, the thrombin detection held a wide linear range from 0.02 pg/mL-20 pg/mL with an R-squared of 0.990 (Fig.4. A-B). Therefore, We can reasonably infer that the design principles and the assay parameters of Captamer fit different aptamer-protein pairs.

To prove that Captamer is independent on the NA/AP sequence, we redesigned a whole new set of Captamer DNA components (Set2) to detect Tau (Fig. 4C). We dissolved Tau in serum to mimic the natural blood sample. Experimental results show that Tau can be rapidly detected in serum down to 2 ng/mL (about 16 pM) which basically meets the diagnostic requirement for Tau concentration of AD patients (10-100pM)^38^.

**Fig. 4.**
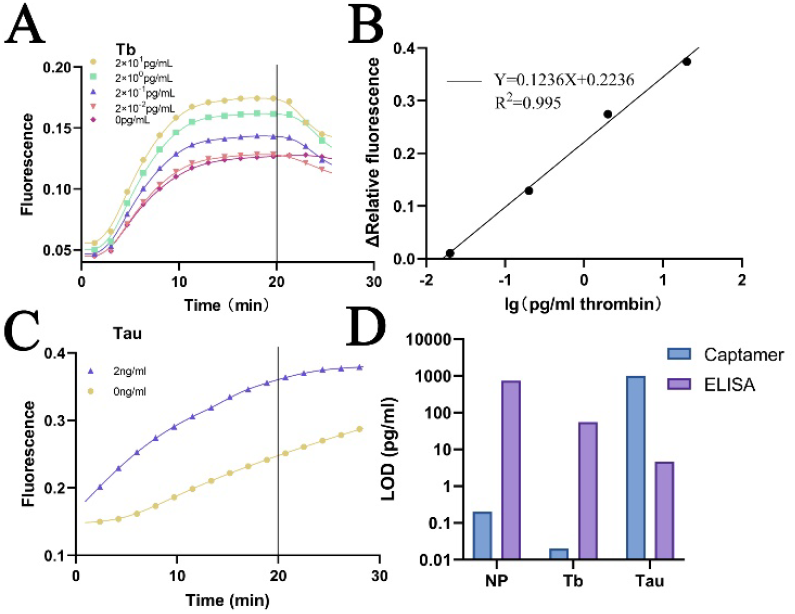
Captamer assays for Tau and Thrombin. **A)** Captamer assay for Tb. **B)** Linear regression fit between the results of the Captamer assay for Tb concentrations (data from A). **C)** Captamer assay for Tau. **D)** Comparison of LOD of Captamer and ELISA.

We conducted a comparison of the limit of detection (LOD) between the captamer assay and the ELISA method for the three target proteins(Fig. 4D). Captamer exhibit significantly higher sensitivity for detecting both NP and Tb. However, the performance of tau is not as good as ELISA. This may be attributed to the lack of affinity of the Tau aptamer employed in the captamer assay.

## DISCUSSION

As an *in vitro* quantitative protein detecting method, Captamer not only manifests a wide detection range, high sensitivity, accuracy, and specificity but also requires little in respect of operating skills and equipment. Taking NP detection as an example, we comprehensively compared the quantitative detecting methods based on aptamers, immunoassays, and electrochemical assays developed in recent years in terms of LOD, detection time (from adding samples to reading results), and preparation time (work prior to adding samples) in Table 2. With the LOD of 0.2 pg/mL, Captamer ranks top among these quantitative protein assays. As for time cost, Captamer is a fast method that only needs 30 min for detection and 1h for the molecular switch preparation. The prepared DNA solution of molecular switch here can be stocked at 4 °C and used for multiple times, so the only time-limiting procedure is the detection, in which all steps can be finished within 30min. In addition, the operations of Captamer assays are just pipetting and mixing solutions and setting a qPCR machine for reaction and fluorescence analysis, which are easy and beginner-friendly. Even people without any professional background can easily master the above operations after a simple training. Besides, Captamer only requires a qPCR machine or, as an cheaper alternative a fluorescence reader as an cheaalternative, both of which are common instrument in biological and clinical laboratories.

**Table 2.**
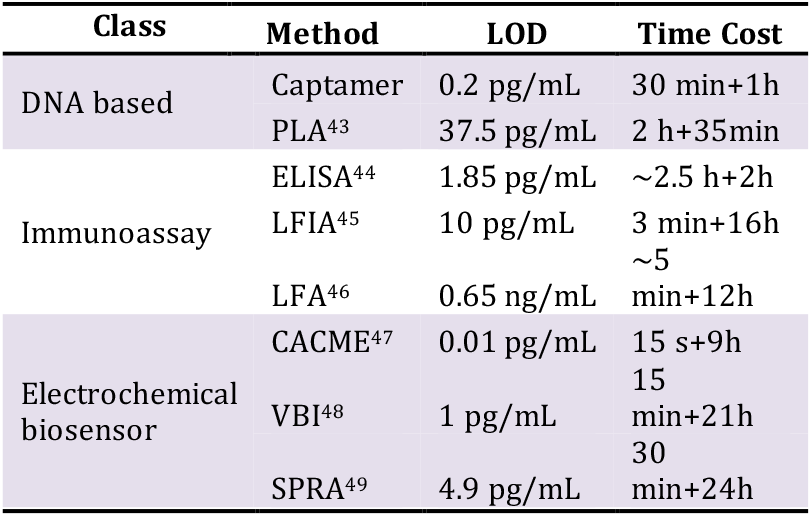
Comprehensive evaluation of NP quantification methods. Time cost means detection time plus preparation time.

We have proved the detectable protein diversity of Captamer with the protein-aptamer pairs of NP, Tb and Tau, suggesting its potential utilization in various protein detection for research or clinic diagnosis like biomarker quantification. When a Captamer system is designed to detect a new target protein, the only change in the original design will be to replace N9-Apt and its binding sequence in the Captamer for NP (Set1) with a high-affinity aptamer of the new protein. It is worth mentioning that most of the current methods for aptamer detection of thrombin are in the pM or even nM range^24,39,40^, and only a few can detect the fM level^41,42^. We performed the thrombin assay without any optimization of the chains, but simply replaced the aptamer, and the detection limit can reach 0.02 pg/mL (about 25 fM), which is already in the forefront of the same type of methods. Set2 used in the Captamer for Tau is an alternative to Set1 in case there is an unexpected sequence similarity between the aptamer and the Set1 DNA sequences, which may lead to incompatibility. The detection sensitivity of the new Captamer will highly depend on the binding affinity of the chosen aptamer. Moreover, Captamer was shown to have good performance in water, saliva, and serum, so it has the potential for detecting proteins in a variety of complex solvent environment in specimen.

Furthermore, Captamer is suitable to be developed as a commercial protein detecting kit. Firstly, except for the reagents from the real-time RPA kit, all other components in Captamer are DNA or DNA derivates, including the molecular switch, two primers, and the probe, which are stable for storage. Secondly, Captamer supports high-throughput screening using 96-well or 384-well plates. Thirdly, Captamer holds a low cost of no more than 6$ per reaction in this study, with an estimated average cost as low as 1$ per reaction under mass production (Supplementary materials). Finally, if the manufacturer prepares and packages the molecular switch solution in the Captamer kit, the assay will be more time-saving (0 min for preparation and 30 min for detection) and user-friendly.

However, Captamer still has some limitations. Firstly, its sensitivity and specificity greatly depend on the aptamer. More specifically, Captamer prefers the aptamer with higher affinity to the target protein and less cross binding to non-target proteins. With the fast development of aptamer screening technologies and the increasing size of the aptamer database, we believe there will be fewer limitations in selecting a suitable aptamer for Captamer in the future. Additionally, Captamer is easily affected by the aerosol contamination of DNA due to its high sensitivity, just like most ultrasensitive DNA detecting methods. Spatially separating the molecular switch preparation and the sample preparation can effectively prevent aerosol contamination.

## CONCLUSION

In this paper, we developed a rapid detection system for microprotein based on nucleic acid aptamer molecular switch and recombinase polymerase reaction, which can be completed in less than 30 min. The detection of SARS-CoV-2 nucleocapsid protein can be done with an accuracy of 0.2 pg/mL, and the results of Tau (1 ng/mL) and thrombin (0.02 pg/mL) shows that our system is universal and has a wide range of potential applications.

## ASSOCIATED CONTENT

This material is available free of charge via the Internet at http://pubs.acs.org. Protocols, Fig. S1-S3, Table S1-S2 (PDF).

## AUTHOR INFORMATION

### Author Contributions

### Notes

The authors declare no competing financial interest.

## ACKNOWLEDGMENT

We sincerely thank the USTC Academic Affairs Office, USTC Department of Life Sciences and Medicine, Core Facility Center for Life Sciences of USTC, and National Training Center for Laboratory Techniques of Life Science of USTC for their tremendous support of this project. Thanks to the University of Science and Technology of China Initiative Foundation for their supports. Thanks to Dr. Jianye Zang, Dr. Xu Li, and Mr. Yongsheng Bai for their enthusiastic help. Thanks to Dr. Haiyan Liu’s lab for providing the instruments. We thank Dr. Zhaofeng Luo for providing suggestions. Thanks to Miss Olivia Wang for her assistance with writing and proofreading. Thanks to the dedication and efforts of other 2021 USTC iGEMers.

## ABBREVIATIONS

RPA: recombinase polymerase amplification
ELISA: enzyme-linked immunosorbent assay
PLA: proximity ligation assay
WB: western blot
LFA: lateral flow assay
MS: mass spectrometer
LAMP: Loop-Mediated Isothermal Amplification
CHA: Catalytic Hairpin Assembly
ESDR: Entropy-driven Strand Displacement Reaction
NAT: PCR-based nucleic acid testing
NP: nucleocapsid protein
Tb: thrombin
Hb: Hemoglobin
THF: Tetrahydrofuran
BE: block efficiency
LFIA: lateral flow immunoassay
CAMEA: capacitive aptasensor coupled with microfluidic enrichment
VBI: voltametric-based immunosensor
SPRA: surface plasmon resonance aptasensor
LOD: limit of detection.

