## Supplemental material for "Captamer: A Novel Quantitative Protein Detecting Method Depending on Aptamer-activated Molecular Switches and RPA Signal Amplification"

### Contents

**Section A：Protocol**

**Section B: How to avoid aerosols**

**Section C: Cost Estimation**

**TableS1: Cost calculation**

**TableS2: DNA Sequences used in this study**

**Figure S1. Asymmetric PCR electrophoresis results**

**Figure S2. Exploration of the optimal temperature**

**Figure S3. MST Results**

### Section A: Protocol

**Asymmetrical PCR:**

The method of asymmetrical PCR is similar to that of normal PCR, only requiring a primer ratio of 20:1. We choose Phanta Max Super-Fidelity DNA Polymerase for its high fidelity and rapid amplification.

*Reaction system

| reagent | volume |
|---|---|
| 2x phanta Max Buffer | 25μl |
| Forward primer (2μM) | 20μl |
| Reverse primer (2μM) | 1μl |
| Template dsDNA (40ng/μl—60ng/μl) | 2μl |
| dNTP Mix (10mM each) | 1μl |
| Phanta Max Super-Fidelity DNA Polymerase | 1μl |

*Reaction process

| Steps | Temperature | Times | Cycles |
|---|---|---|---|
| Initial Denaturation | 95°C | 3min | — |
| Denaturation | 95°C | 15sec | 30 |
| Primer Annealing | 57°C | 15sec | 30 |
| Extension | 72°C | 15sec | 30 |
| Final Extension | 72°C | 5min | — |

**Electrophoresis:**

1. Prepared 2% agarose gel. In fact, the separation of single and double strains was better at higher concentrations of agarose gel. However, considering that the gel extraction steps would be complicated if the gel concentration is higher than 2%. We choose the 2% agarose gel after comprehensive consideration.

2. Electrophoresis was performed at 120V for 60 min. (ssDNA moves faster than dsDNA)

3. After electrophoresis, excise the ssDNA from the agarose gel. The excised gel should be as thin as possible.

**ssDNA Gel Extraction:**

We use QIAEX® II Gel Extraction Kit (QIAGEN) for gel recovery and conduct the gel extraction as the manual.

1. Excise the DNA band from the agarose gel with a clean, sharp scalpel. Use a 1.5 ml microfuge tube for processing up to 250 mg agarose per tube.

2. Weight the gel slice in a colorless tube, Add Buffer QX1 according to DNA fragment size: 6 volumes for <100bp; 3 volumes for 100bp-4kb;3 volumes with 2 volumes of water for >4kb. Add 6 volumes of Buffer QX1 when using >2% of Metaphor agarose gels.

3. Resuspend QIAEXⅡ by vortexing for 30s. Add QIAEXⅡto the sample and mix: Use 10 μl QIAEXⅡ for ≤2μ g DNA； 30 μl for 2-10 μg DNA； and an

additional 30 μl for each additional 10 μg DNA.

4. Incubate at 50 °C for 10 min to solubilize the agarose and bind the DNA, Mix the sample by vortexing every 2 min to keep QIAEXⅡ in suspension. Check that the color of the mixture is yellow. If the color of the mixture is orange or violet, add 10 μl 3 M sodium acetate, pH 5.0, and mix. The incubation should then be continued for at least 5 min.

5. Centrifuge the sample for 30 s and carefully remove supernatant with a pipet.

6. Wash the pellet with 500 μl Buffer QX1. Resuspend the pellet by vortexing. Centrifuge the sample for 30 s and remove all traces of supernatant with a pipet. This wash step removes residual agarose contaminants.

7. Wash the pellet twice with 500 μl Buffer PE. Resuspend the pellet by vortex, Centrifuge the sample for 30 s and carefully remove all traces of supernatant with a pipet. This step removes residual salt contaminants.

8. Air-dry the pellet for 10-15 min or until the pellet becomes whites. If 30 μl QIAEXⅡ suspension is used, air-dry the pellet for approximately 30 min. Do not vacuum dry, as over drying, may lead to decreased elution efficiency.

9. To elute DNA, add 20 μl of 20 mM Tris-Cl, TE buffer (10 mM Tris-Cl, 1 mM EDTA, pH 8.0) or water and resuspend the pellet by vortex. Incubate according to the DNA fragment size: 5 min at room temperature (15-25°C) for ≤4kb; 5 min at 50°C for 4-10 kb; or 10 min at 50°C for >10kb.

10. Centrifuge for 30s, and carefully pipet the supernatant into a clean tube. The supernatant now contains the purified DNA.

11, Optional: repeat steps 9 and 10 and combine the eluates. A second elution step will increase the yield by approximately 10-15%.

Notes:

*The protocol is for cleanup of DNA fragments of 40bp to 50kb.

*A heating block or water bath at 50°C is required.

*All centrifugation steps are carried out at 17900 x g (~13000 rpm) in a conventional tabletop microcentrifuge at room temperature (15-25°C).

**Construction of molecular switch**

1. Calculate the concentration of ssDNA obtained from the gel extraction. In order to obtain the accurate concentration of single chain, a piece of empty gel of similar size should be extracted according to the same steps. With the empty gel as blank control, the actual spectrum of ssDNA can be obtained by subtracting the measured value from the blank control. Use a Microsoft Excel Application to predict the spectrum of single strand. Compared the actual value of 260/230 and 260/280 with those calculated by program. If the two value is close, the ssDNA can be regarded as pure. Predict the theoretical absorbance of corresponding single DNA strand (1M). Use the $A_{260}$ of the single DNA chain to subtract the $A_{260}$ of empty gel from the measured single chain to obtain the actual single chain concentration.

$$c_{ssDNA} = \frac{A_{260,ssDNA} - A_{260,empty}}{A_{260,\mathrm{predictivevalue}}}$$

2. Add the collected ssDNA, aptamer and $ddH_2O$ to the PCR tube.

3. A series of temperature gradients are set by PCR amplifier as follows:

| | | | | | | |
|---|---|---|---|---|---|---|
| **Temperature** | 95°C | 89°C | 83°C | 77°C | 71°C | 65°C |
| **Time** | 5min | 5min | 5min | 5min | 5min | 5min |
| **Temperature** | 59°C | 53°C | 47°C | 41°C | 35°C | 29°C |
| **Time** | 5min | 5min | 5min | 5min | 5min | 5min |

**RPA**

DNA Thermostat Rapid Fluorescence Kit (Nanjing warbio Biotechnology Co., Ltd) was used for RPA.

1. Add 11.5 μL $ddH_2O$ and 2 μL nucleic acid template into the tube caps successively. The volume of nucleic acid template added can be adjusted according to the concentration of nucleic acid, and the volume of $ddH_2O$ added can be adjusted accordingly. Similarly, protein solutions can be added and the volume of $ddH_2O$ should be adjusted, so that the total volume of the template, protein solution and $ddH_2O$ is 13.5 μL

2. Add 2.5 μL B Buffer into the tube caps.

3. Add 2 μL upstream primer (10 μM), 2 μL downstream primer (10 μM) and 0.6 μL probe (10 μM) into the cap of the reaction tube.

4. Add 29.4 μL A buffer into each enzyme powder tube. After enzyme powder completely dissolute, transfer all the solution into an EP tube. Fully mix and reload the solution into the reaction tubes.

5. Fasten the reaction tube on the tube cap. Flip the reaction tube upside down for 8~10 times, and centrifuge quickly.

6. Immediately put the reaction tube into the fluorescence detection equipment at 37°C and collect the FAM channel every 30 seconds.

Notes:

*Reaction system

| Ingredient | Volume(μL) |
| --- | --- |
| A buffer | 29.4 |
| Forward Primer(10μM) | 2 |
| Reverse Primer(10μM) | 2 |
| Probe (10μM) | 0.6 |
| dd$H_2O$, DNA template and protein solution | 13.5 |
| B buffer | 2.5 |
| Total volume | 50 |

*Due to the high sensitivity of the kit, please pay attention to avoid nucleic acid contamination during the reaction and set a blank control.

*A buffer needs to be thoroughly melted and mixed.

**Microscale Thermophoresis：**

We use the Monolith His-Tag Labeling Kit RED-tris-NTA 2nd Generation to achieve fluorescent labeling of protein. The affinity of N protein and aptamer was detected by MST.

Step A: Affinity of dye to target protein

Preparation：Prepare empty PCR-tubes 1-16 for gradient dilution.

1. Suspend the dye in 25 μL of buffer to obtain 5 μM dye solution;

2. Prepare 200 μL of 50 nM solution of the RED-tris-NTA 2nd Generation dye in buffer by mixing 2 μL of dye (5 μM) and 198 μL buffer;

3. Prepare 30 μL of 4 μM His-tagged protein in buffer;

4. Transfer 10 μL buffer into PCR-tubes 2-16;

5. Transfer 20 μL of 4 μM His-tagged protein solution into the first PCR-tube;

6. Transfer 10 μL of the ligand from PCR-tube 1 to PCR-tube 2 with a pipette and mix by pipetting up-and-down multiple times. Transfer 10 μL to tube 3 and mix. Repeat the procedure for tubes 4-16. Discard the extra 10 μL from tube 16;

7. Add 10μL of 50 nM RED-tris-NTA 2nd Generation dye solution to each tube (1-16) and mix by pipetting;

8. Incubate for 30 min at room temperature;

9. Load the capillaries and measure the samples at 40% percent excitation power and medium MST power.

*To ensure a high labeling interest, we recommended using PBS or alternatively HEPES buffer and a pH in the range of 7-8 for the labeling reaction. As the affinity between the dye and the His-tag decreases significantly in Tris buffers and at a pH below 7, these conditions are not advised.

If the affinity of the RED-tris-NTA 2nd Generation dye to the His-tagged protein of interest is stronger than 10 nM ($K_d$≤10nM), please continue with step B.

Step B: Protein labeling

1. Adjust the protein concentration to 200 nM in a volume of 100 μL;
2. Mix 90 μL of protein (200 nM) with 90 μL of dye (100 nM);
3. Incubate for 30 min at room temperature;
4. Centrifuge the sample for 10 min at 4°C and 15000×g and transfer the supernatant to a fresh tube;
5. The protein is labeled and ready for the binding assay;

Step C: Binding assay

1. Prepare 25 μL of the ligand (aptamer) at 2× concentration in the assay buffer. Make sure to avoid buffer mismatches within your titration series;
2. Add 10 μL of buffer into the PCR-tubes 2-16;
3. Transfer 20 μL of the ligand into PCR-tube 1;
4. Transfer 10 μL of the ligand from PCR-tube 1 to PCR-tube 2 with a pipette and mix by pipetting up-and-down multiple times. Transfer 10 μL to PCR-tube 3 and mix. Repeat the procedure for PCR-tube 4-16. Discard the extra 10 μL from PCR-tube 16;
5. Add 10 μL of labeled protein to each tube (1-16) and mix by pipetting. The

final target protein concentration in the assay is 50 nM. This concentration should be used for the calculation of the $K_d$ value;

6. Load the capillaries and measure the samples. Recommended settings are 40% excitation power and medium MST power. At the final dye concentration of 25 nM the expected fluorescence intensity at 40% excitation power is around 300 counts on a Monolith NT.115.

### Section B: How to avoid aerosols

1. The laboratory needs to have negative or positive pressure isolation conditions such as a fume hood or ultra-clean bench to prevent aerosols from entering the sample. If your lab or work environment is negative pressure, perform double-stranded dilutions in it, which will prevent double-stranded molecules from entering the template sample. If your lab or work environment is positive pressure, perform single-strand preparation, molecular switch assembly in it.
2. Use a gun with a filter membrane. Through a lot of experiments we found that the main factor causing laboratory aerosol contamination is that the pipette gun will adsorb some aerosol molecules, which can be well avoided by the filter membrane.
3. Open the centrifuge tube or PCR tube with DNA solution as gently as possible.
4. Dispense and freeze all reagents used. Replace all aerosols once they are found to be contaminated. The cost of checking where the contamination is much higher than replacing the reagents.

### Section C Cost Estimation

1. The RPA reaction kit provides 48 reactions. Since the kit is less popular in China, the production cost is still high. If it can be produced on a large scale, the production cost can be reduced significantly.
2. It seems that the probe with four modifications are expensive. However, the amount purchased once is 100 μM for 100 μl, which first needs to be diluted to 10 μM, and only 0.6 μl for each reaction, so one purchase can be used 1666 times.
3. The aptamer also has a blocking group modification, and again, it is 100 μM for 100 μl at the time of purchase. The aptamer works at a concentration of 1 pM, so a gradient dilution is required for use. It can be used 100000 times with one purchase.
4. The primers are 100 μM for 100 μl at the beginning of the purchase, and their working requirement is 10 μM for 2 μl, so they can be used 500 times in one purchase.

**TableS1: Cost Calculation**

| Projects | Price $ | Average single response $ |
|---|---|---|
| RPA reaction kit | 239.55 | 4.99 |
| Probe | 261.33 | 0.16 |
| Aptamer | 116.15 | 0.00 |
| Primers | 4.65 | 0.01 |
| Total | | 5.16 |

**Table S2：DNA Sequences used in this study**

| | | |
|---|---|---|
| **NP** | **Positive Control** | acatacagccaagcgttaaccctaactcggtattgcgctggatgtgtcaatgtagcggtgcccctaaggaatattgtcgtaagcgacatccagctaaatctttccaagcatcgttcagatgatctacagaatgcatcgagcccctttcaatattaactgccaactcactttgagtgtttaacactgtatcgctactgtcattaggtactaccgaggcaatatccgcgcctcgagtcgtcagtccttctgtgatctctat |
| | **CoVIIN-NA dsDNA** | acatacagccaagcgttaaccctaactcggtattgcgctggatgtgtcaatgtagcggtgcccctaaggaatattgtcgtaagcgacatccagctaaatctttccaagcatcgttcagatgatctacagaatgcatcgagcccctttcaacctcgagtcgtcagtccttctgtgatctctat |
| | **N-NA-primer1** | atagagatcacagaaggactgacgac |
| | **N-NA-primer2** | acatacagccaagcgttaaccctaactcgg |
| | **AP dsDNA** | taaatctttccaagcatcgttcagatgatctacagaatgcatcgagcccctttcaatattaactgccaactcactttgagtgtttaacactgtatcgctactgtcattaggtactaccgaggcaatatccgcgcctcgagtcgtcagtccttctgtgatctctat |
| | **AP-primer1** | taaatctttccaagcatcgttcagatgatctac |
| | **AP-primer2** | atagagatcacagaaggactgacgac |

| | | |
|---|---|---|
| | **Nor-apt** | gctggatgtcgcttacgacaatattccttaggggcaccgctacattgacacatc cagc[C3-spacer] |
| | **N9-apt** | cgcttacgacaatattccttaggggcaccgctacattgacacatccagc[C3-spacer] |
| | **Probe** | agtagcgatacagtgttaaacactcaaag[FAM-dT][THF]ag[BHQ-dT]tggcagttaata[C3spacer] |
| | **Primer1** | cgcggatattgcctcggtagtacctaatgac |
| | **Primer2** | acatacagccaagcgttaaccctaactcgg |
| **Tau441** | **T-NA dsDNA** | gctgacgcaacttacgctcttcgtgaggaaacgtcagtttaatacttttactggtg ctgaagaagaaaagcctcgtcaaagacctgacaggaatccatcgatatctcc acattccttcagattcctccatgcatacacgaaggtatgccatcgcctgtgaag ccgaagtcaagttcgaaggtgacaccttggtgaacagaagactg |
| | **T-NA-primer-1** | gctgacgcaacttacgctcttcgtgaggaaacg |
| | **T-NA-primer-2** | cagtcttctgttcaccaaggtgtcaccttcg |
| | **T-AP dsDNA** | aatccatcgatatctccacattccttcagattcctccatgcatacacgaaggtat gccatcgcctgtgaaggaatcatgagttccgatgatgctgtcgcgccgaagtc aagttcgaaggtgacaccttggtgaacagaa |
| | **T-AP-primer-1** | aatccatcgatatctccacattccttc |

| | | |
|---|---|---|
| | **T-AP-primer-2** | ttctgttcaccaaggtgtcacc |
| | **Aptamer** | cctgtcaggtctttgacgaggcttttcttc[C3spacer] |
| | **Probe** | gaatttcagaggctatagcgatctcagg[FAM-dT]a[THF]a[BHQ-dT]cgatagatcgcta[C3spacer] |
| | **Primer1** | gctgacgcaacttacgctcttcgtgaggaaacg |
| | **Primer2** | gcgacagcatcatcggaactcatgattcc |
| **Thrombin** | **Tb-NA dsDNA** | acatacagccaagcgttaaccctaactcggtattgcagtcaccccaacctgccctaccacggactttaaatctttccaagcatcgttcagatgatctacagaatgcatcgagcccctttcaacctcgagtcgtcagtccttctgtgatctctat |
| | **Tb-NA-primer-1** | atagagatcacagaaggactgacgac |
| | **Tb-NA-primer-2** | acatacagccaagcgttaaccctaactcgg |
| | **AP dsDNA** | taaatctttccaagcatcgttcagatgatctacagaatgcatcgagcccctttcaatattaactgccaactcactttgagtgtttaacactgtatcgctactgtcattaggtactaccgaggcaatatccgcgcctcgagtcgtcagtccttctgtgatctctat |
| | **AP-primer-1** | taaatctttccaagcatcgttcagatgatctac |
| | **AP-primer-2** | atagagatcacagaaggactgacgac |

| | |
|---|---|
| **Aptamer** | agtccgtggtagggcaggttggggtgact [C3spacer] |
| **Probe** | agtagcgatacagtgttaaacactcaaag[FAM-dT][THF]ag[BHQ-dT]tggcagttaata[C3spacer] |
| **Primer1** | cgcggatattgcctcggtagtacctaatgac |
| **Primer2** | acatacagccaagcgttaaccctaactcgg |

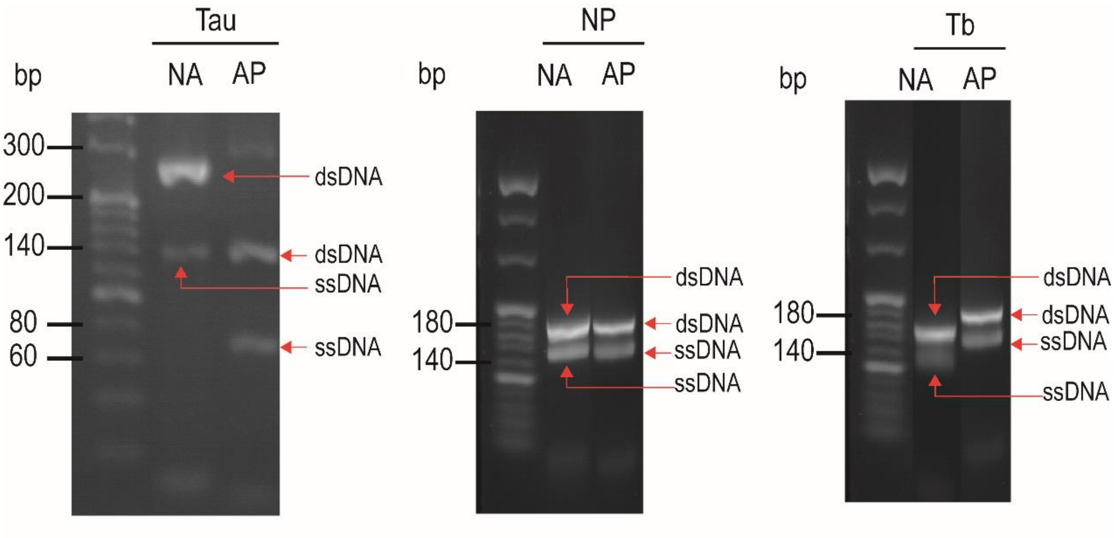


**Fig. S1：Agarose Gel Electrophoresis of Asymmetric PCR preparation for DNA single-stranded.** The left figure shows NA and AP chains for Tau detection, the middle shows NA and AP chains for NP detection and the right shows NA and AP chains for Tb detection. For those three pictures, the strand on the left are NA chains and the right are AP chains. The single strand bands are all below the double stands. T-NA dsDNA size is 212bp while T-AP dsDNA is 139bp. N-NA dsDNA size is 182bp and N-AP dsDNA is 168bp. Tb-NA dsDNA size is 154bp while Tb-AP dsDNA is 168bp.

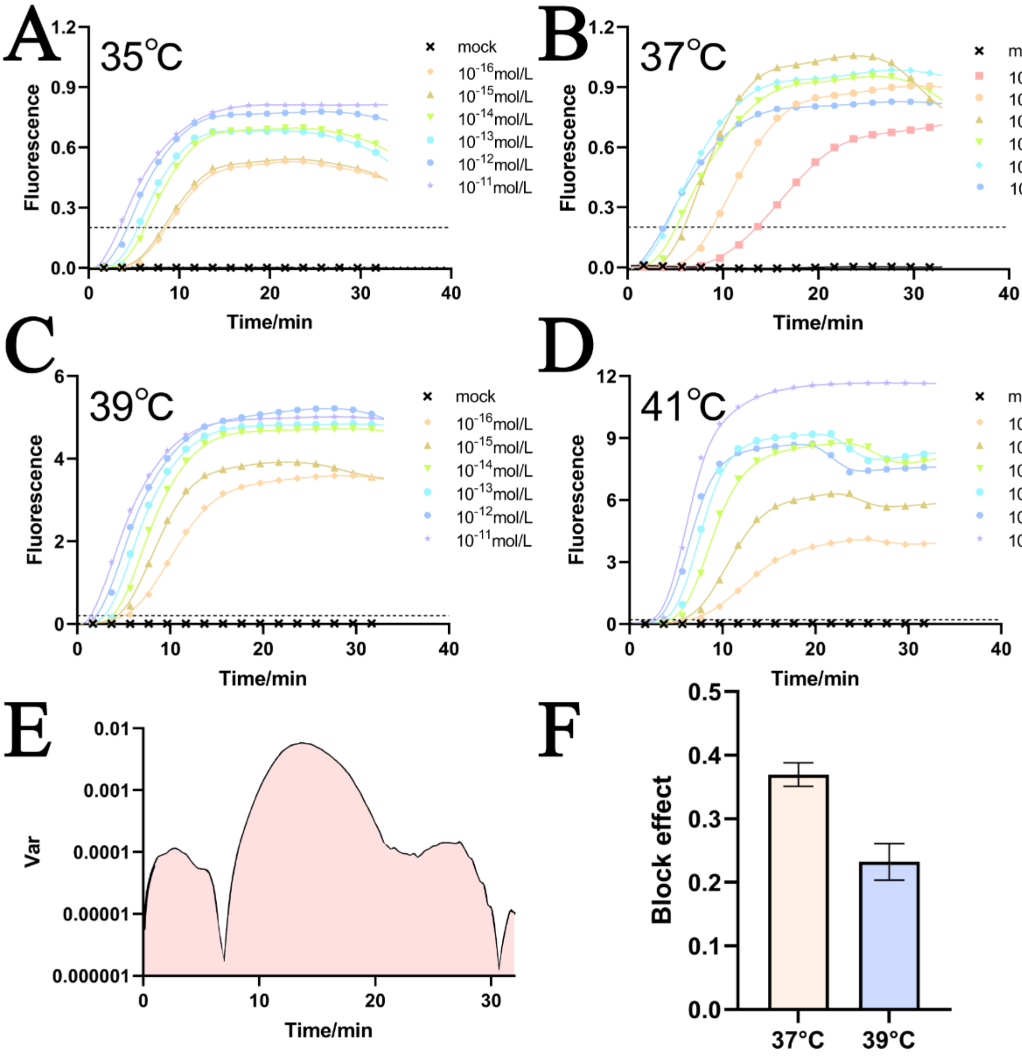


**Fig. S2：Temperature gradient. A-D：35°C, 37°C,39°C, 41°C Fluorescence curve.** E： Analysis of variance. F. Comparison of BE values at 37°C and 39°C. We combined the curve fluorescence intensity magnitude, discrete nature and deterrence effect, and finally chose 37°C as the optimal reaction temperature.

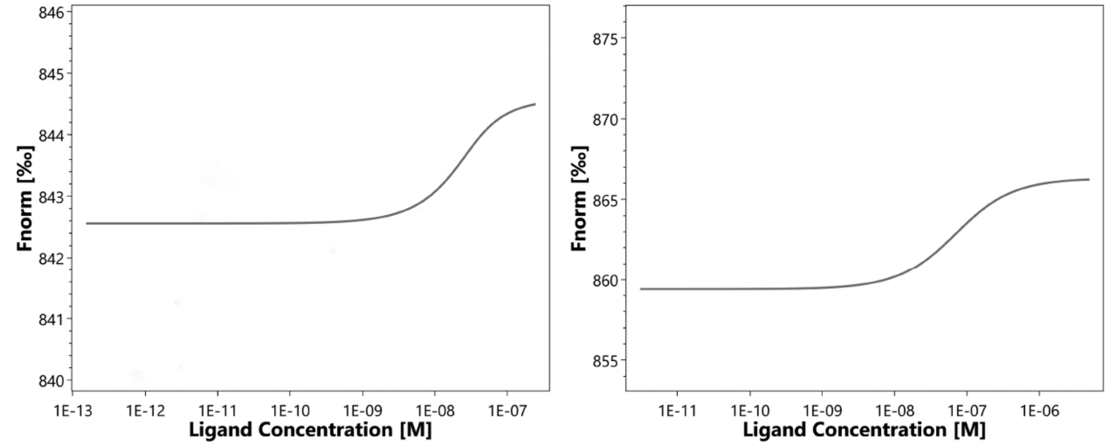


**Fig. S3: MST Affinity Curve Analysis. Left: N9-Apt Right: Nor-Apt.**
